# Exploring the relationship between dolphins and fisheries: uncovering the spatial and temporal patterns that influence potential conflicts along Portugal’s north coast

**DOI:** 10.64898/2026.03.25.714190

**Authors:** Beatriz Barbieri, Luís Afonso, Cláudia Oliveira-Rodrigues, Iolanda Silva, Ágatha Gil, Ana Marçalo, Isabel Sousa-Pinto, Ana Mafalda Correia, Raul Valente

## Abstract

The north coast of mainland Portugal supports a strong dolphin presence and extensive fishing activity, increasing the likelihood of interactions, such as bycatch. This study provides an initial assessment of potential conflict areas, using automatic identification system (AIS) data from Global Fishing Watch. To this end, sighting data from the ATLANTIDA project (2021–2024) on the common dolphin (*Delphinus delphis*) were used to describe spatiotemporal patterns of occurrence and encounter rates, and to predict their association with fishing effort to identify and map areas of potential overlap. A generalised additive model (GAM) was then applied, integrating environmental, spatial, temporal, and fisheries-related variables to identify the main predictors of species occurrence. Common dolphins were frequently observed during the summer, with an average encounter rate of 3.662 sightings/km. This high encounter rate may be associated with factors such as sea surface temperature, diet, and purse seine fishing activity. The maps showed a spatial overlap between fishing grounds and areas of common dolphin occurrence. Fishing effort was nearly identical between locations with sightings (2.00 h/km²) and those without (1.62 h/km²), suggesting that dolphins are not actively avoiding fishing areas but may instead frequent them due to shared habitat preferences. The best-fitted GAM indicated that encounters were related to year, latitude, fishing effort, depth, sea surface temperature, and season. There was an increase in occurrence over the years and a decrease with increasing fishing effort and sea surface temperature, possibly linked to changes in prey availability, although broad confidence intervals warrant cautious interpretation. Despite some limitations encountered in this study, we believe our findings provide valuable insights into the relationship between dolphin occurrence, environmental conditions, and fishing activities in the area, establishing an important baseline for future conservation and fisheries management efforts.

## Introduction

Interactions between cetaceans and worldwide fisheries are likely to occur, as fisheries activities often overlap with cetacean distribution (Kiszka et al., 2022). As a result of this kind of interaction, cetaceans and fishing gear may come into direct physical contact, which could cause unintentional capture, also known as bycatch (Dias et al., 2022). Bycatch, even in small-scale fisheries, can result in serious injuries or death (Alexandre et al., 2022) and is largely believed to be the primary cause of extinction risk in small cetaceans (Temple et al., 2024).

The Iberian Atlantic waters, located in the Eastern North Atlantic Ocean, are home to a variety of cetacean species (Goetz et al., 2014; Marçalo et al., 2024). The high productivity and rich marine resources of the Iberian Atlantic waters not only attract cetaceans by providing abundant prey and suitable habitats, but also support extensive fishing activity, heavily exploited by Spanish and Portuguese fisheries (Goetz et al., 2014). Fisheries in the area, especially purse seining, target various species that overlap significantly with the preferred prey of local cetacean populations (Goetz et al., 2014), such as sardine (*Sardina pilchardus*), horse mackerel (*Trachurus hurus*) or anchovy (*Engraulis encrasicolus*) (Marçalo et al., 2018). This overlap results in a degree of competition between cetaceans and fisheries for shared resources (Goetz et al., 2014).

Among the cetacean species found in the Atlantic waters of the Iberian Peninsula, the short-beaked common dolphin (*Delphinus delphis*) is the most prevalent (Marçalo et al., 2024). Despite their abundance (Paradell et al., 2019), this species is classified as “Nearly Threatened (NT)” in Mainland Portugal (Ferreira et al., 2023), and interactions with fisheries could lead to a local population decline over time (Paradell et al., 2019). Along the Portuguese mainland coast, bottom-set nets, including trammel and gill nets, are thought to be the gear most likely to cause these interactions, with common dolphins being the most affected (Silva & Sequeira, 2003; Goetz et al., 2014; Vingada & Eira, 2018; Marçalo et al., 2024). Likewise, some studies conducted along Portugal’s northern coast (e.g. Dias et al., 2022; Marçalo et al., 2025; Wise et al., 2007) revealed that common dolphins were frequently observed during purse seine fishing operations. Portuguese purse seine fisheries primarily target sardines, a significant prey species for common dolphins (Dias et al., 2022; Marçalo et al., 2025). They operate mostly from sunset to sunrise, which aligns with the dolphins’ feeding period and increases the chance of interactions (Wise et al., 2007).

To prevent this from escalating further, it is important to analyse the temporal and spatial patterns of common dolphins’ presence during fishing activities, as well as the impact of fishing efforts on cetaceans’ presence (Marçalo et al., 2015). Hence, the present work aimed to provide a comprehensive analysis of cetacean-fishery interactions through the identification of bycatch risk areas for common dolphins on the northern Portuguese coast.

## Materials and Methods

### Study Area

The present study was conducted along the northern coast of mainland Portugal, from Espinho to Caminha, at distances ranging from 2 to 12 nautical miles from the coastline (40.984–41.798°N, 9.141–8.677°W) (Figure 1). The north Portuguese shelf is characterised by a broad continental shelf (40-70 km) (Dias et al., 2022) and has a relatively uniform topography and coastline (Vitorino et al., 2002). The shelf dynamics in this area are frequently influenced by the seasonal patterns of two major atmospheric systems, the Azores High and Icelandic Low (Vitorino et al., 2002). The pressure gradient between the Azores High and the Iberian Peninsula, along with a thermal low over central Iberia, drives northerly winds, sustaining an upwelling regime from July to September with favourable conditions (Vitorino et al., 2002). This supports high marine biodiversity, including many economically significant species (Veiga-Malta et al., 2019; Zwolinski et al., 2010), making fishing a culturally significant and traditional practice in this area (Mendes et al., 2019).

**Figure 1.**
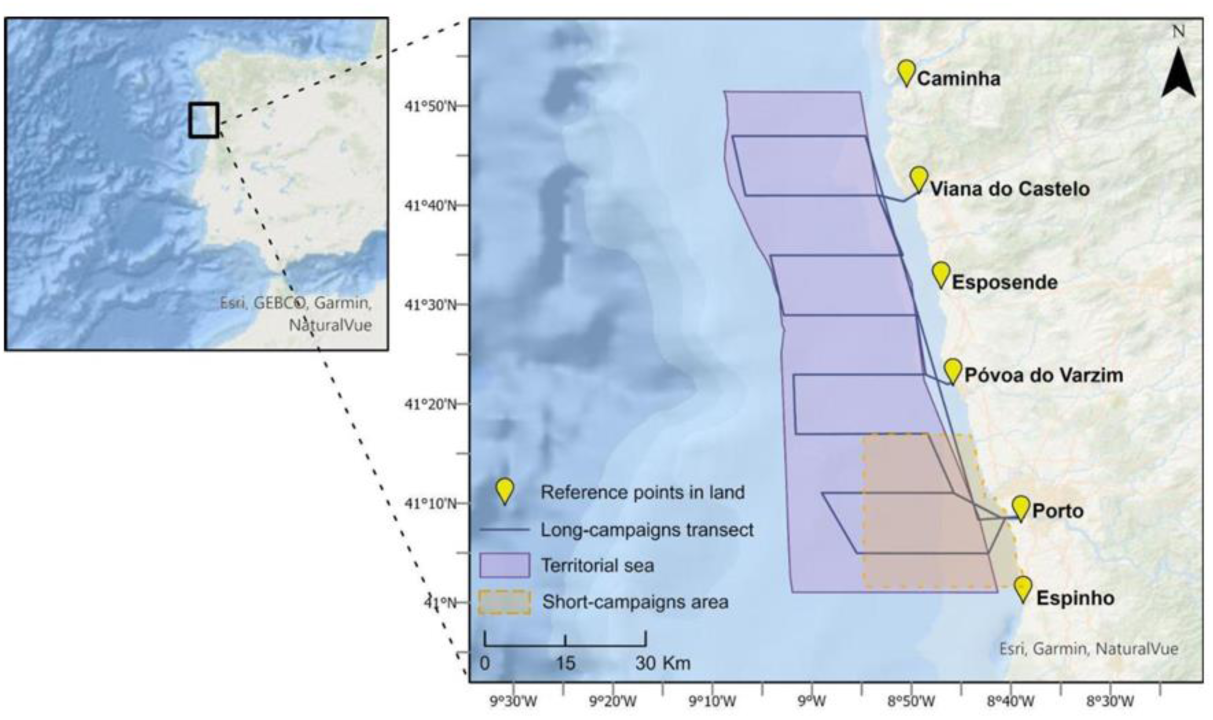
Study area (Espinho to Caminha), with respective monitoring campaign transects and in-land reference points (Retrieved from Afonso, 2023).

### Data Collection and Monitoring

Sighting data from the ATLANTIDA project, conducted from 2021 to 2024, were collected using two different sampling strategies: (1) during long campaigns – conducted twice a year, one in summer (July-September) and one in winter (January-March), each lasting four days along predefined transects (Figure 1); (2) during short campaigns – conducted throughout the year, consisting of four-hour monitoring sessions carried out along random transects (Figure 1). In both campaign types, visual monitoring was conducted using binoculars (magnification of 7 ×50 mm, with scale and compass), following a detailed observer protocol adapted from Correia et al. (2015). The applications “Locus Map” (used from 25/06/2021 to 14/07/2023) and “ILogWhales” (used from 21/07/2023 to 23/04/2024) were operated on a tablet with GPS to record effort transect coordinates and register sightings and other sampling locations. Survey effort and data were systematically recorded, including species identification, group size, and observed behaviour. When a precise count of individuals was not possible, the minimum and maximum number of animals was recorded, along with the observer’s estimated most likely number of individuals (best estimate). Survey periods are classified as either “on-effort” (active surveying) or “off-effort” (interrupted surveying, e.g. whenever a dedicated 180° survey effort cannot be maintained or conditions are unfavourable for cetacean monitoring). Only “on-effort” data was considered to ensure standardised sampling effort and to minimise detection bias. For additional information on the sampling protocol, see Correia et al. (2015).

Monthly fishing activity data were sourced from the Automatic Identification System (AIS) records provided by Global Fishing Watch (GFW, https://globalfishingwatch.org/). This approach has become increasingly common in scientific research focused on cetaceans and serves as a valuable tool for addressing interactions between fisheries and cetaceans (Moore et al., 2021; Tack, 2025). Global Fishing Watch analyses AIS data from vessels identified as commercial fishing and applies an algorithm that classifies each transmission as “apparently fishing” or “not fishing”, based on changes in speed and direction (Global Fishing Watch, 2024). Throughout this study, fishing activity was obtained from a tabular dataset in which apparent fishing events were aggregated into a predefined grid, producing the total apparent fishing effort per cell and linking it to the active vessels operating in or near each location (Hintzen et al., 2025).

Environmental variables considered in this study included static factors, such as depth, and dynamic oceanographic parameters, including sea surface temperature (SST), chlorophyll-a concentration (CHL), and sea surface salinity (SSS). Gridded bathymetry data (depth) were retrieved from the General Bathymetric Chart of the Oceans (GEBCO, https://www.gebco.net/) database for the entire study area. SST, CHL, and SSS were obtained from the Copernicus Marine Environmental Monitoring Service (CMEMS) platform (https://marine.copernicus.eu/), using the MGET (Marine Geospatial Ecology Tools) toolbox (Roberts et al., 2010) from ArcGIS Pro 3.5 (Esri, 2025). For more details on the spatial and temporal resolutions, as well as temporal coverage for each variable, see table in the supplementary section.

## Data Analysis

### Encounter Rate Analysis

On effort sightings of common dolphins were recorded during all monitoring campaigns, alongside survey effort. Encounter rates for each season were then calculated using the following expression:

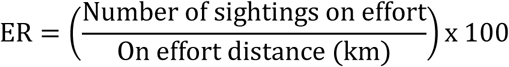

Where the “number of sightings on effort” comprises only those sightings made during active surveys, and the “on effort distance (km)” refers to the total distance travelled, in kilometres, while actively surveying for cetaceans, as recorded by GPS. Distances were calculated in ArcGIS Pro 3.5 (Esri, 2025) using the WGS 1984 Geographic Coordinate System (EPSG:4326), projected to World Mercator (EPSG:3395).

### Spatial and Temporal Analysis

To assess potential interactions between common dolphins and fishing activities, a heat map of fishing effort for the entire monitoring period was produced across our study area, alongside sightings of the study species. Additionally, seasonal maps based on spatial grids were developed to analyse the spatial and temporal patterns of common dolphin presence during fishing activity. For each season of the year, three types of grids were created:

i) Fishing Effort (FE) grid – composed of binary values (0 and 1), where 0 corresponds to cells with FE values below the seasonal average, and 1 corresponds to values above the seasonal average.
ii) Encounter Rate (ER) grid – also composed of binary values (0 and 1), where 0 represents cells with ER values below the seasonal average, and 1 represents values above the seasonal average.
iii) Overlap grid between FE and ER – resulting from the combination of the two previous grids, classified into three categories (0, 1, and 2), where 0 indicates that both ER and FE are below average, 1 indicates that at least one of the parameters

is above average, and 2 indicates that both are above average. This grid represents a gradient of potential overlap, corresponding respectively to a low, medium and high probability of interaction between cetaceans and fishing activities.

All spatial analyses and map production were carried out in ArcGIS Pro 3.5 (Esri, 2025).

### Ecological Modelling Analysis

Data distributions were visually assessed (e.g., Q–Q plots and density plots) and tested using the Shapiro-Wilk test. Statistical modelling was then used to assess the non-linear relationship between the occurrence of common dolphins and a set of environmental predictors, as well as temporal, spatial, and fisheries-related variables. To this end, Generalised Additive Models (GAMs) were chosen. GAMs offer a more flexible model by using smooth terms (Hastie & Tibshirani, 1986; Wood, 2003), and they have been widely used to explain cetacean distribution (Brotons et al., 2004; Pearce & Boyce, 2006; Correia et al., 2015, 2019). As explanatory variables, we selected one topographic variable (depth), three oceanographic variables (SST, CHL, and SSS), two spatial variables (longitude and latitude), three temporal variables (month, season, and year), and one fisheries-related variable (fishing effort). The encounter rate was used as the response variable. Additionally, all GAMs were performed using the *mgcv* 1.9.4 package (Wood, 2004, 2011) in R 4.5.0. (R Core Team, 2023). Before modelling, collinearity among explanatory variables was assessed using Pearson correlation, with a threshold of 0.75 (Correia et al., 2019, 2021; Marubini et al., 2009). Longitude and depth were the only set of variables highly correlated. Moreover, multicollinearity involving multiple variables was evaluated using the Variance Inflation Factor (VIF), with a threshold of 3 (Correia et al., 2019, 2021; Zuur et al., 2010). All VIF values fell below the established threshold, so no additional variables needed to be eliminated. Considering the greater ecological relevance of depth relative to longitude, the depth variable was retained, and a model with a negative binomial distribution (with a log link function) was fitted (Correia et al., 2020). Moreover, a maximum of four splines (k-fold set to 4) were used to limit the complexity of the smoothers, modelling the effects of the explanatory variables (Correia et al., 2019, 2021). To identify the best-fitting model, a saturated binomial GAM was fitted to assess their association with the occurrence of common dolphins, followed by a backward selection (Correia et al., 2015, 2019, 2020, 2021). After backward selection, the best final GAM model included the following explanatory variables: year, season, latitude, fishing effort, sea surface temperature, and depth.

The best models were selected by using the Akaike Information Criterion (AIC, Burnham & Anderson, 2002). The model with the lowest AIC was chosen at each step, and when AIC differences were less than 2, models were compared using a Chi-squared test (Zuur et al., 2007).

Whenever differences between AIC values were not statistically significant (p-value > 0.05), the simpler model was preferred (Burnham & Anderson, 2002). Additionally, when a spline term exhibited an estimated degree of freedom close to 1, indicating near-linearity, the smooth term was replaced with a linear function (Correia et al., 2021). The final model was then evaluated using simulated residuals employed in the *DHARMa* 0.4.7 package, together with the “gam.check” function and multiple diagnostic plots obtained from the “appraise ()” function (*gratia* 0.10.0 package).

## Results

### Encounter Rate Analysis

Throughout the three-year monitoring period, a total of 51 survey days were conducted, consisting of 25 short and 26 long campaigns. During these campaigns, a total of 108 common dolphin sightings on effort were recorded throughout the entire study area. The total on-effort distance (km), number of sightings, and encounter rate (sightings/km) were calculated for each season of the campaign – winter (January-March), spring (April-June), summer (July-September), and autumn (October-December). The total surveyed distance amounted to 4,423.191 km, with the highest recorded in summer (1832.500 km) and the lowest in autumn (345.085 km). During winter, the total on-effort distance corresponded to 1446.123 km, whereas in spring it was 799.483 km. Sightings peaked in summer (64 sightings) and had the lowest record in autumn (7 sightings). Winter and spring accounted for 20 and 17 sightings, respectively. The highest encounter rate was recorded in summer (3.662 sightings/km), followed by spring (2.022 sightings/km), autumn (2.017 sightings/km) and winter (1.413 sightings/km).

### Spatial and Temporal Analysis

Fishing effort data along the northern coast of mainland Portugal were combined with common dolphin sighting locations to analyse spatial patterns of co-occurrence. The resulting map (Figure 2) revealed that areas of fishing activity overlapped with dolphin hotspots, particularly near the coast. On average, monthly apparent fishing hours were 1.62 h/km^2^ in non-sighting and 2.00 h/km^2^ in sighting areas. Despite the relatively similar mean values between sighting and non-sighting areas, slightly higher fishing activity was observed in sighting areas.

**Figure 2.**
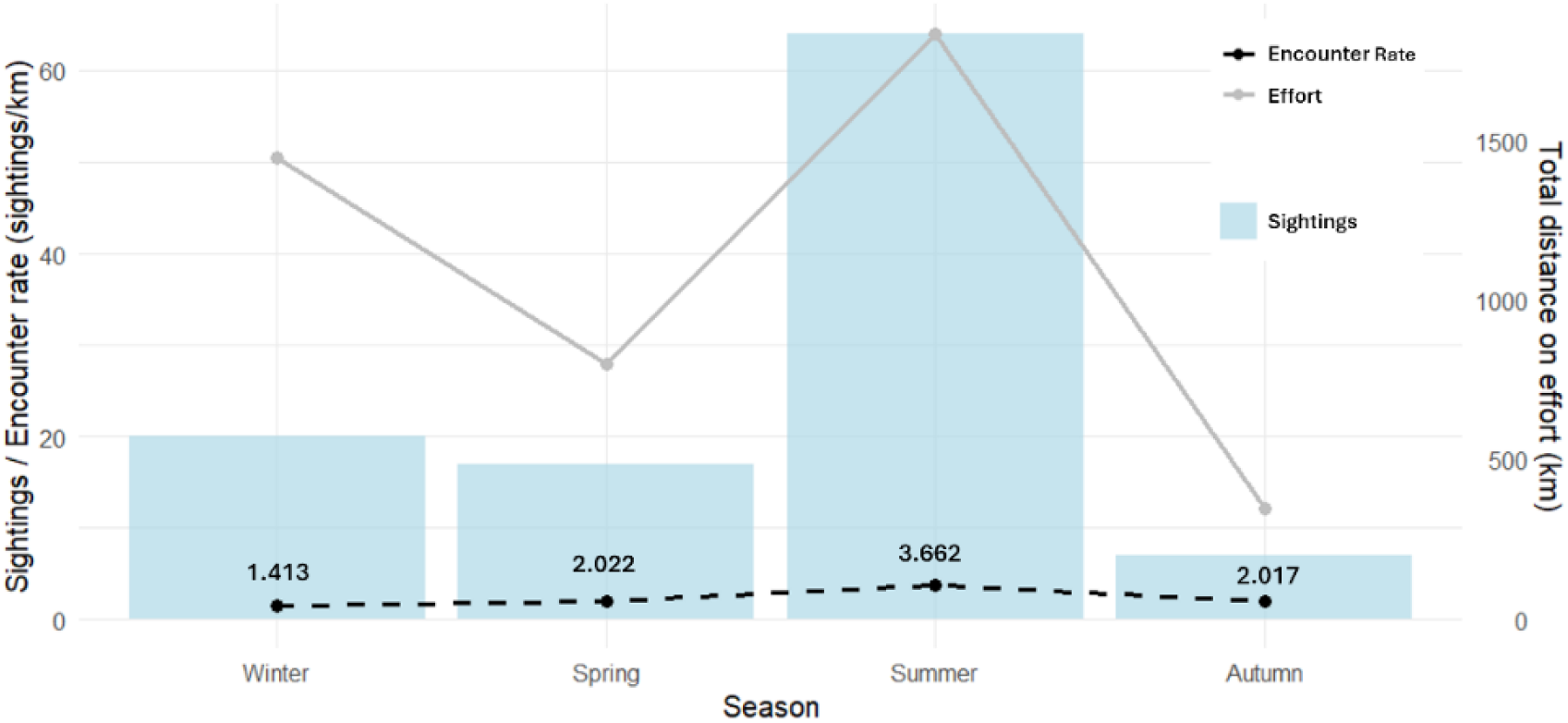
Common dolphin (*Delphinus delphis*) occurrence, encounter rate (sightings/km), and survey effort (km) per season, considering only on-effort sightings. Seasonal encounter rate values correspond to the mean calculated across all years, whereas survey effort and the number of sightings were obtained by summing all records within each season.

Analysis of grid-based maps showed a balance between cells with fishing effort values that were below and above the average across all seasons (Figure 3).

**Figure 3.**
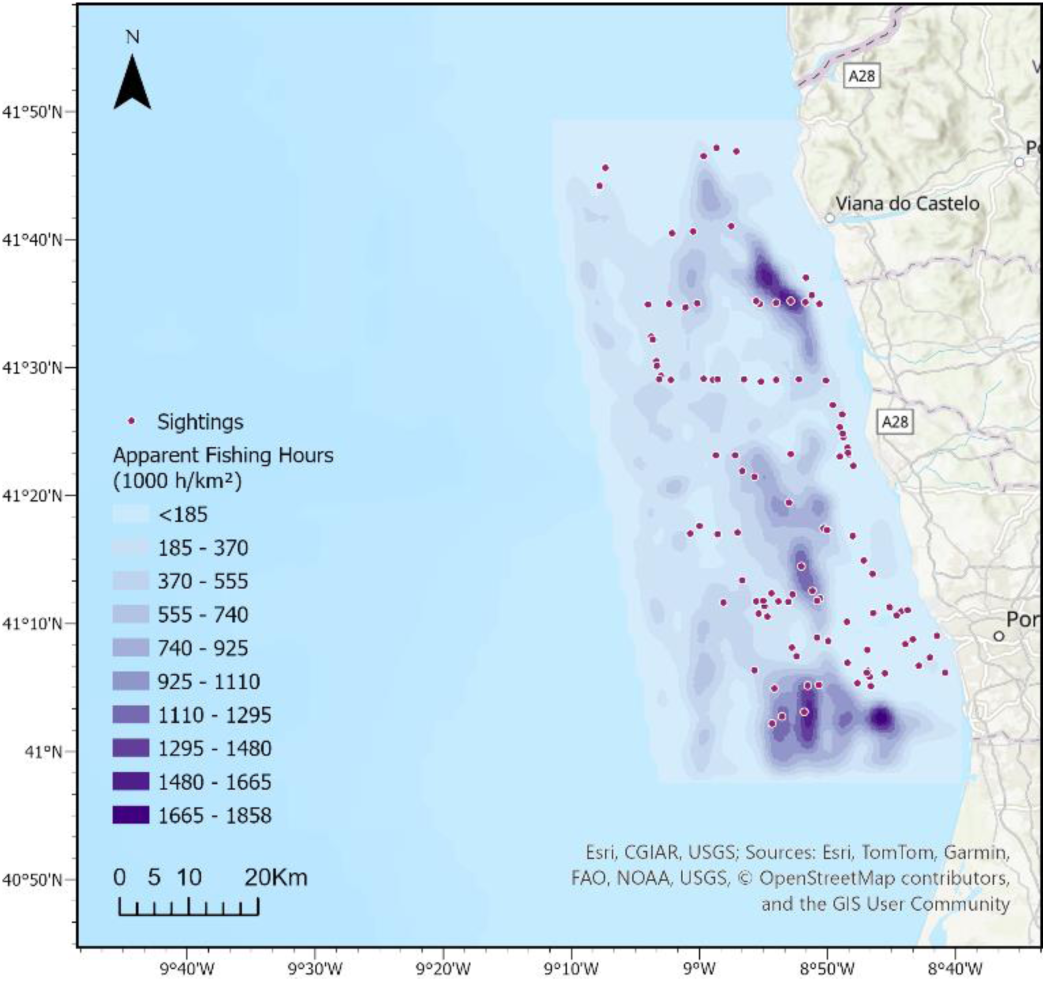
Spatial variation of common dolphin (*Delphinus delphis*) occurrence and fishing activity (by apparent fishing hours (h/km^2^)) along the northern coast of mainland Portugal. Only on-effort sightings were considered. Red dots represent individual dolphin sightings, while the purple colour gradient reflects the total apparent fishing hours in each specific area on the day of the sightings.

Additionally, a similar, relatively stable spatial pattern was observed over the years, suggesting that the distribution of fishing effort remains consistent throughout all seasons. Regarding the encounter rate, a greater discrepancy was observed in the distribution of values, with a predominance of cells with values below the average. Furthermore, it was not possible to identify a clear and consistent spatial pattern between seasons. By examining the correlation between fishing effort and encounter rate, overlap cells were identified in all seasons, corresponding to areas where both variables exceeded their respective averages. Summer had the highest number of overlapping cells (13), followed by winter (7), spring (4) and autumn (1). This suggests that there is a higher risk of interaction during the summer months.

### Ecological Modelling Analysis

The variables retained in the best final GAM model are indicated in Table 1. GAM results indicated that the Summer and Winter seasons, as well as SST, Year, and Lat had p-values < 0.05, indicating statistically significant differences between the encounter rate and these predictors. CHL and SSS were not retained in the final model.

**Table 1.** Results from the best Generalised Additive Model (GAM). In bold are the p-values < 0.05, indicating statistically significant differences. Lat – latitude. SST – sea surface temperature.

| Best GAM |  |  |  |  |
| --- | --- | --- | --- | --- |
| Encounter_Rate ~ s(Year) + Season + s(Lat) + s(Fishing_Effort) + SST + s(Depth) |  |  |  |  |
| Variables | Estimate | Standard Error | Z-Value | Pr(>\|z\|) |
| Intercept | 6.3361 | 1.9968 | 3.173 | **0.00154\*** |
| SeasonSpring | - 0.5750 | 1.1327 | -0.508 | 0.61178 |
| SeasonSummer | 2.9143 | 0.9643 | 3.022 | **0.00255\*** |
| SeasonWinter | -2.5364 | 1.2096 | -2.097 | **0.03617\*** |
| SST | -0.3047 | 0.1152 | -2.644 | **0.00829\*** |
| Variables | Degrees of Freedom | Reference Degrees of Freedom | Chi-Square | P-value |
| s(Year) | 1.968 | 2.379 | 7.029 | **0.000584\*** |
| s(Lat) | 2.841 | 2.980 | 4.320 | **0.004493\*** |
| s(Fishing_Effort) | 1.911 | 2.243 | 1.154 | 0.336297 |
| s(Depth) | 2.634 | 2.901 | 2.081 | 0.064671 |

By observing the fitted GAM plots, it is clear that the encounter rate of common dolphins increased over the years (Figure 4). Additionally, the encounter rate fluctuates across the latitudinal range of the study area, peaking at 41.0°N, decreasing, increasing again at 41.3°N, and finally decreasing once more at 41.5°N. Regarding fishing effort, a descending trend can be observed in relation to dolphin encounters in the area. However, the large confidence intervals limit the strength of this interference. Furthermore, depth showed a similar oscillating trend to latitude, with an increase in the encounter rate between values of −60 and −30. In addition, dolphin encounters were found to decrease with increasing sea surface temperature values. Nonetheless, broad confidence intervals indicate that this trend should be interpreted cautiously. Finally, the frequency of encounters appeared to be higher during the summer, followed by fall, spring and winter.

**Figure 4.**
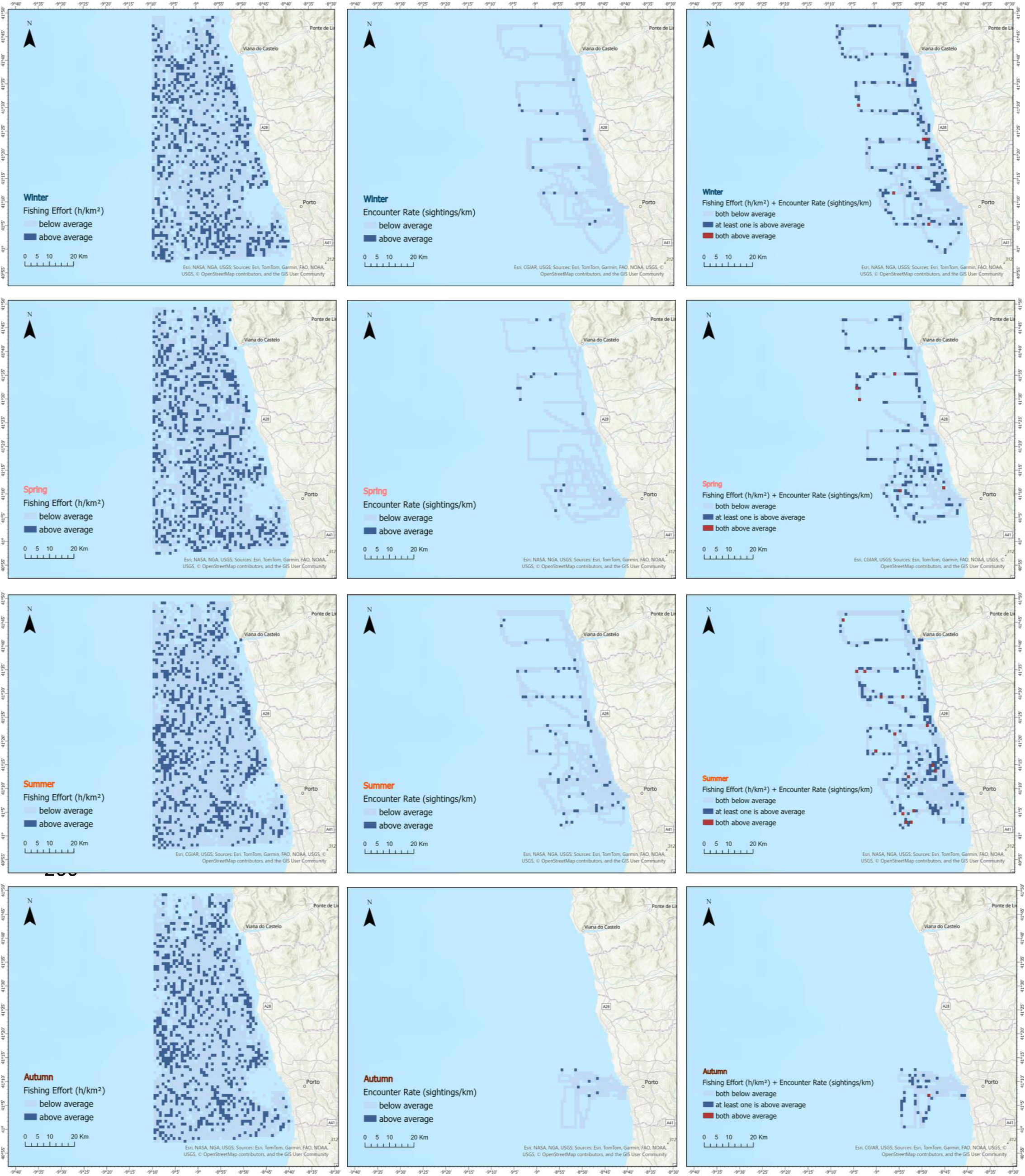
Gridded maps showing the spatial and temporal patterns of fishing effort (left), encounter rate (middle), and overlap hotspots between fishing effort and encounter rate (right) across the study area for each season. The resulting grid-based maps were visualised using graduated colour symbology, with each colour representing a distinct class of fishing effort, encounter rate, or degree of overlap: grey indicates below-average values, blue indicates above-average values, and red represents grid cells where both fishing effort and encounter rate are above average (overlap).

**Figure 5.**
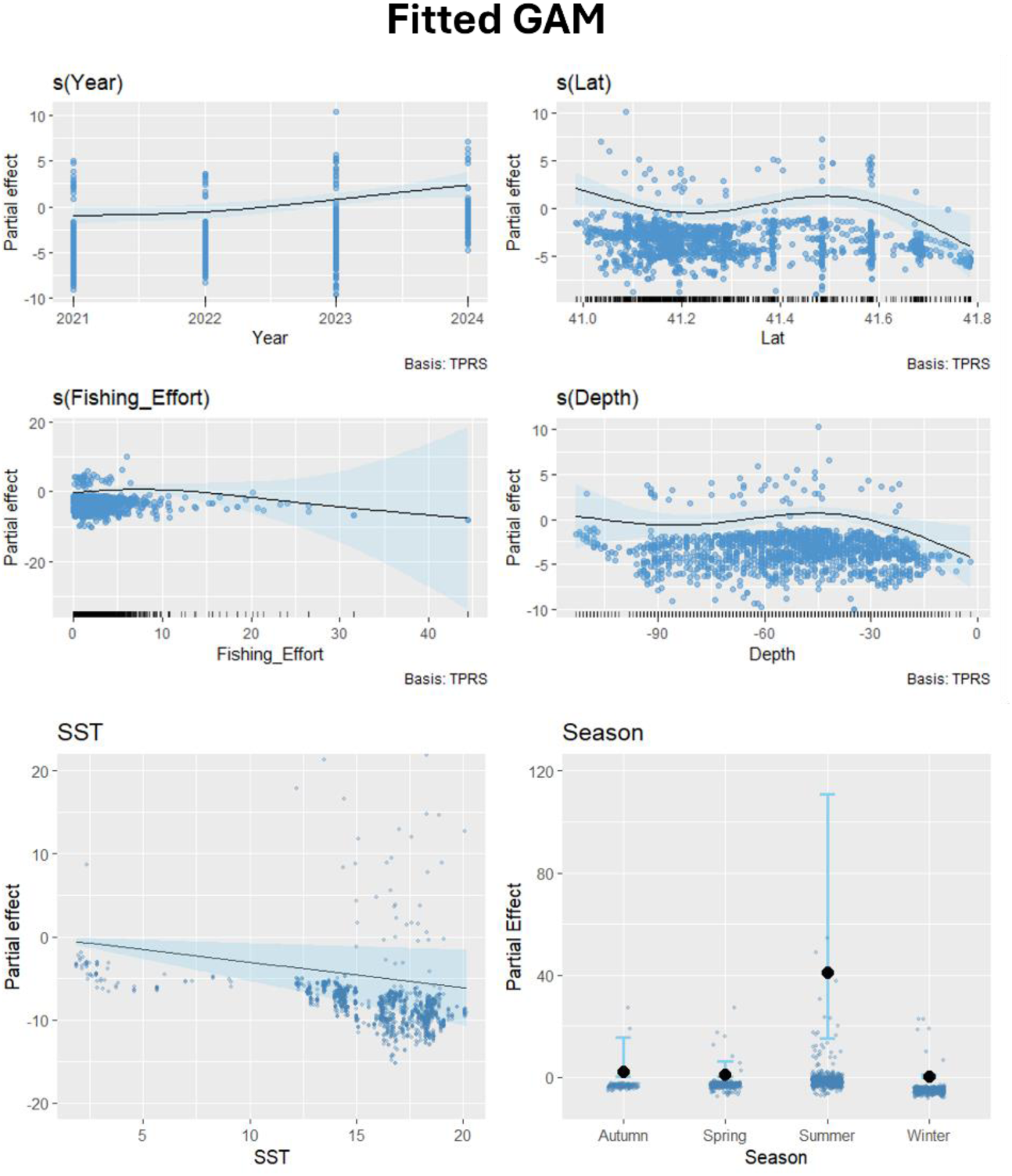
Partial effect plots of the final Generalised Additive Model (GAM) predicting the probability of encounter of common dolphins (*Delphinus delphis*) in the study area. The blue shadow around the linear predictor represents the 95% confidence intervals of the spline functions, while blue dots within the graph area correspond to the residuals. Lat – latitude. SST – sea surface temperature.

Kolmogorov–Smirnov test did not indicate significant deviations of the residuals from the expected distribution (p > 0.05). However, the dispersion test indicated a significant deviation (p < 0.05), suggesting that the variance of the data is not fully captured by the model. Additionally, the outlier test was not significant (p > 0.05), and the graph of residuals versus predicted values showed no significant problems detected, suggesting an adequate overall fit.

Additionally, there are no significant influential data points (with “hat” values below 0.25), and the model residuals do not follow a normal distribution.

## Discussion

### Encounter Rate Analysis

The prevalence of the common dolphin in our research area was expected, as it is the most abundant cetacean throughout the Western Iberian Peninsula (Gilles et al., 2023; Paradell et al., 2019). Moreover, previous studies on the occurrence and distribution of cetaceans within the Portuguese Continental Exclusive Economic Zone (EEZ) have consistently demonstrated a strong prevalence of this species along the northern coast, with an apparent preference for northern latitudes compared to the rest of the mainland (Brito & Sousa, 2011; Correia et al., 2015; Gil, 2017; Gilles et al., 2023, Brouder et al., 2025). Additionally, common dolphins were observed every year of monitoring and in every season. This was expected, as the literature reports year-round occurrence of the species along the Portuguese coast (Bencatel et al., 2019; Brouder et al., 2025; Correia et al., 2015; Gil, 2017; Moura et al., 2012). Summer recorded the highest number of sightings, which corresponds to the season with the greatest survey effort. This pattern is consistent with the positive relationship between survey effort and sightings reported by Correia et al. (2020) and may partly explain the low number of sightings observed in autumn, when effort was lowest. However, this pattern does not apply to the encounter rate. Although summer simultaneously presents the highest sampling effort and the highest encounter rate, the lowest value did not occur in autumn, but rather in winter. This may be due to unfavourable weather conditions during this period, which may have reduced the probability of detection. On the other hand, there may also be a high sampling effort relative to the number of individuals present in the area, which reaches a “plateau”. Once this point is achieved, high survey efforts will no longer increase the number of records, as most individuals have already been detected. This indicates that survey effort alone does not fully explain the true seasonal pattern of common dolphin occurrence and that other factors should be considered. For instance, sea surface temperature may play a role, as Fernández et al. (2013) reported a preference of common dolphins for warmer waters within the northwest Iberian Peninsula. In addition, seasonal shifts in prey availability may further contribute to the observed pattern. The sardine, the preferred prey of the common dolphin (Silva, 1999; Marçalo et al., 2018), exemplifies the typical life strategy of small pelagic fish: a short lifespan, rapid growth, high reproductive capacity, and an extended spawning period (Santos et al., 2012). The spawning of sardines off Portugal occurs predominantly between November and April, with peak activity along the northwest coast mainly during winter (Ré et al., 1990; Chícaro et al., 2003; Santos et al., 2012). Thus, this seasonal dynamic may help explain the peak in common dolphin occurrence in summer, when prey availability increases after spawning. Furthermore, Portuguese purse seine activity intensifies from late spring to early autumn, driven by market demand for target species (Feijó, 2013; Marçalo et al., 2015; Feijó et al., 2018). So, dolphins may approach coastal areas to benefit from increased sardine availability and prey aggregations created by purse seine operations (Marçalo et al., 2015; Dias et al., 2022; Brouder et al., 2025), showing a higher occurrence during this period.

### Spatial and Temporal Analysis

Our results also revealed a spatial overlap between fishing grounds and areas of common dolphin occurrence. Although the fishing effort values between non-sighting and sighting areas were very similar, averages were slightly higher in areas where dolphins were sighted, suggesting that common dolphins may prefer habitats that overlap with fishing grounds. Notably, several studies revealed that common dolphins might find it beneficial to feed on prey schools that are brought to the surface or discarded during commercial fishing activities (e.g. Silva, 1999; Cruz et al., 2016, 2018). This suggests the hypothesis that interactions in our study area are likely to occur. Indeed, dedicated studies aimed at identifying cetacean–fishery interaction hotspots have shown that the northwest coast (Wise et al., 2007; Marçalo et al., 2015; Dias et al., 2022), particularly between 40°N and 42°N (Dias et al., 2022), is the area with the highest probability of interactions involving common dolphins, especially with purse seining (Wise et al., 2007; Dias et al., 2022). This is mostly due to the mutual interest in the same resource: sardines (Silva, 1999; Marçalo et al., 2018; Dias et al., 2022). The northern region of Portugal operates the most purse seines and accounts for the highest number of sardine catches (Feijó, 2013; Feijó et al., 2018), thereby increasing the chances of encounters in this area (Dias et al., 2022).

While total landings have changed over time, Portuguese fisheries exhibit stable temporal trends and generally consistent spatial patterns in landings and assemblage structures across regions (Sousa et al., 2005; Bueno-Pardo et al., 2020), which aligns with our findings. Fishing seasons remained relatively consistent throughout the study period, likely because they are linked to the species’ spawning or recruitment periods (Bueno-Pardo et al., 2020). The encounter rate, however, shows greater variability, with more values below the average. This is likely because surveys frequently covered areas without sightings. Furthermore, even when dolphins are spotted, a high survey effort typically results in a lower overall encounter rate. Additionally, there was a noticeable overlap of cells exhibiting high fishing activity and encounter rates in all seasons. Summer had the greatest number of overlapping cells, suggesting it is the season with the highest likelihood of interactions. This observation aligns with findings by Marçalo et al. (2015), who noted that interactions with common dolphins were more frequent in summer, when the demand for target species increases.

### Ecological Modelling Analysis

Generalised Additive Model (GAM) results indicated that the encounter rate of common dolphins increased over the years and showed statistically significant differences. Following the implementation of the Iberian Management Plan (2011–present), which severely restricted catches, the sardine stock began to show signs of recovery (Silva et al., 2015; ICES, 2023), and in recent years (2016–2024), the spawning biomass appears to have been increasing (Monteiro et al., 2025). This recovery may have contributed to a gradual return of the common dolphin to more coastal areas to feed, reflecting the observed increase in encounter rates. This pattern is consistent with studies conducted on minke whales (*Balaenoptera acutorostrata*) on the northwest coast of the Iberian Peninsula, where it was reported that, after a period of dietary diversification during the decline of sardines, the diet returned to being dominated by this species post-stock recovery (Monteiro et al., 2025). Additionally, Marçalo et al. (2015) noted that fish abundance is also influenced by latitude, which may help explain the significant effect detected for this variable. Nevertheless, this result should be interpreted cautiously due to the relatively narrow latitudinal range of our study area, which limits the detection of potential spatial preferences.

Our results further suggest a tendency for common dolphin encounters to decrease with increasing fishing effort, although this relationship was not statistically significant. This aligns with findings from Marçalo et al. (2015), who observed that when sardine catches were higher, the likelihood of cetacean presence increased at lower levels of fishing effort. However, the absence of a significant relationship indicates that increased fishing activity does not appear to influence dolphins’ presence in the area in a way that would prevent spatial interactions. Instead, it suggests that these areas may have lower prey availability, leading fisheries to spend more time searching for resources (Dias et al., 2022), which may also influence dolphin distribution. Additionally, a high fishing effort value may also reflect the combined activities of multiple vessels operating from the same port (Dias et al., 2022).

Furthermore, depth was effectively retained in the model, as several studies consistently indicate that depth is a reliable predictor of common dolphin presence (e.g. Cañadas et al., 2002; Cañadas & Hammond, 2008; Moura et al., 2012; Correia et al., 2015, 2019). However, it did not show a significant effect on common dolphin occurrence. This may be related to the relatively narrow depth range within the study area, which is largely restricted to the shallow continental shelf. Although the GAM results suggest that dolphin occurrence remains relatively stable with increasing depth and shows slightly lower values at depths below approximately 30 metres, depth may not represent a strong environmental gradient influencing dolphin distribution.

Furthermore, an increase in sea surface temperature negatively impacts the presence of common dolphins. As mentioned by Correia et al. (2015), sea surface temperature (SST) serves as an effective indicator of upwelling events, which bring cold, nutrient-rich waters to the surface (Caldeira et al., 2002; Mason, 2009). This may clarify the observed preference for cooler waters, as common dolphins favour more productive regions associated with strong upwellings (Cañadas & Hammond, 2008; Moura et al., 2012; Correia et al., 2015; Halicka, 2016). Seasonal analysis revealed significant differences in encounters, with summer showing the highest rates, followed by fall, spring, and winter. The same trend was observed by Spadoni et al. (2025), who explained that water temperature and chlorophyll content may play a role. The summer upwelling season from June to September positively affects the recruitment of small pelagic species, such as sardines and horse mackerel (Santos et al., 2014; Spadoni et al., 2025), influencing the habitat suitability for common dolphins, with higher values observed during this season (Spadoni et al., 2025).

Although chlorophyll and sea surface salinity have been described to influence common dolphins’ distribution (Forney, 2000; Cañadas & Hammond, 2008; Moura et al., 2012; Halicka, 2016; Correia et al., 2019; Paradell et al., 2019), they were not retained in the final GAM in the present study. This suggests that these variables may not have significantly enhanced model performance or accounted for additional variation in encounter rates. This may indicate that, within the environmental range covered by our study area, these variables may not strongly influence dolphin distribution.

### Limitations and Future Perspectives

Despite these results, this study has some limitations that should be considered. First, there is always the possibility of not detecting dolphins even when they are present, as they are animals that spend most of their time underwater (Evans & Hammond, 2004; Paradell et al., 2019; Marçalo et al., 2024). To overcome the limitations of visual surveys, passive acoustic monitoring and environmental DNA (eDNA) could be implemented, allowing detections at night or when animals are submerged.

Additionally, fishing effort data were obtained from AIS rather than collected in real time during dolphin observations. GFW offers many benefits, as it is open to the public, and data can be collected for various regions worldwide (Wardhani et al., 2022). However, AIS has inherent limitations that restrict its coverage mainly to mid to large-scale vessels fishing in offshore waters (≥ 12 m) (Wardhani et al., 2022). Small-scale fisheries (< 12 m) are not required to have a device installed on board that allows automatic location and identification of the vessel through the vessel monitoring system (Regulation (EC) No 1224/2009, Article 9). Although recent laws require all EU vessels, without exception, to register and report their catches digitally, some small fishing vessels may be exempt from this obligation until 2030, and all small-scale fleets will have up to four years to adapt to these new requirements. Consequently, AIS data often fail to capture vessels that are relevant in our study area, and that may lead to interactions with common dolphins. Moreover, Wardhani et al. (2022) reported that mapping the distribution of AIS-tracked fishing vessels at a higher resolution compromises the detection of interannual seasonal signals due to low absolute vessel presence observed in each grid cell. In addition, Rufino et al. (2023) discovered that certain trips were related to routes where fishers were merely moving around or just verifying if the area contained the needed resources, rather than actually fishing. Hence, misinterpretation of fishing effort results in inaccurate assessments of the impact of fishing on various ecosystem aspects (Hintzen et al., 2025), as well as the impact on common dolphins. Moreover, although spatial overlap between fisheries and dolphins was identified, this does not necessarily indicate that interactions or bycatch occurred, as no direct bycatch data were available to validate this assumption. Similarly, it’s important to note that cells classified as “no overlap” do not necessarily represent a real absence of potential interactions with the common dolphin. The high proportion of cells with encounter rates below the average may reflect areas where no sightings were recorded, rather than an actual absence of the species. Consequently, this can lead to an underestimation of their occurrence, particularly in areas with higher risk. So, a way to overcome both the limitations of AIS and the misinterpretation of overlapping risk zones would be the collection of real-time fishing activity data, ideally in collaboration with fishers, as this would improve the accuracy of overlap estimates compared to effort data derived from global platforms. In parallel, incorporating bycatch records from observer programs or even through reports from fishers is essential to verify whether spatial overlap with fishing gear translates into actual interactions or mortality events. Future research could also integrate stranding network data with drift modelling approaches to improve the identification of areas with higher bycatch risk.

Additionally, the oceanographic variables used in the analysis were sourced from remote sensing products instead of in situ measurements, which may not capture detailed variability. In a limited study area such as this one, data with low spatial resolution represent only a subset of the environmental drivers that may influence the common dolphin’s distribution.

## Conclusion

Despite these limitations, our findings offer valuable insights into the relationship between dolphin occurrences, environmental conditions, and fishing activities in the area. This research enhances our understanding of potential interactions between common dolphins and fisheries along the northern coast of Portugal. Ultimately, these results reinforce the need for mitigation efforts that account for both ecological and socio-economic dimensions, promoting coexistence between fisheries and cetaceans.

## Acknowledgements

This study was partially supported by the CETUS project and occurrence data collected under the ATLANTIDA project (2021–2024). We thank all team members and collaborators involved in data collection and support throughout the study period.

## Funding information

This work is a result of the project ATLANTIDA II (ref. NORTE2030-FEDER-01799200), co-financed by the European Union through the NORTE 2030 Regional Program and the European Regional Development Fund (ERDF). The research study was conducted within the framework of the REDUCE project (GA: 101135583) funded by the European Union through the Horizon Europe programme. Ph.D. fellowships for authors LA (2024.04444.BD), CR (2023.02122.BD), and AG (PD/BD/150603/2020) were supported by Portuguese Foundation for Science and Technology (FCT).

## Notes

### Competing Interest Statement

The authors have declared no competing interest.

## References

Afonso, L. (2023). Environmental DNA: a novel cetacean monitoring technique in the north coast of Continental Portugal. Master’s Thesis. University of Aveiro.

Alexandre, S., Marçalo, A., Marques, T. A., Pires, A., Rangel, M., Ressurreição, A., Monteiro, P., Erzini, K., & Gonçalves, J. M. (2022). Interactions between air-breathing marine megafauna and artisanal fisheries in Southern Iberian Atlantic waters: Results from an interview survey to fishers. Fisheries Research, 254. 10.1016/j.fishres.2022.106430

Bearzi, G., Bonizzoni, S., & Gonzalvo, J. (2011). Dolphins and coastal fisheries within a marine protected area: Mismatch between dolphin occurrence and reported depredation. Aquatic Conservation: Marine and Freshwater Ecosystems, 21(3), 261–267. 10.1002/aqc.1179

Bencatel, J., Sabino-Marques, H., Álvares, F., Moura, A. E., & Márcia Barbosa, A. (2019). *Atlas de Mamíferos de Portugal*.

Brito, C., & Sousa, A. (2011). The environmental history of cetaceans in portugal: Ten centuries of whale and dolphin records. In PLoS ONE (Vol. 6, Number 9). 10.1371/journal.pone.0023951

Brotons, L., Thuiller, W., Araújo, M. B., & Hirzel, A. H. (2004). Presence-absence versus presence-only modelling methods for predicting bird habitat suitability. Ecography, 27(4), 437–448. 10.1111/j.0906-7590.2004.03764.x

Brouder, S., Marques, T. A., Oliveira, N., Monteiro, P., Gonçalves, J. M. S., & Marçalo, A. (2025). When Sardines Disappear: Tracking Common Dolphin, *Delphinus delphis*, Distribution Responses Along the Western Iberian Coast. Animals, 15(11). 10.3390/ani15111552

Bueno-Pardo, J., Pierce, G. J., Cabecinha, E., Grilo, C., Assis, J., Valavanis, V., Pita, C., Dubert, J., Leitão, F., & Queiroga, H. (2020). Trends and drivers of marine fish landings in Portugal since its entrance in the European Union. ICES Journal of Marine Science, 77(3), 988–1001. 10.1093/icesjms/fsaa010

Burnham, K., & Anderson, D. (2002). Model Selection and Multimodel Inference: a Practical Information-theoretic Approach. (2nd ed). Springer.

Caldeira, R. M. A., Groom, S., Miller, P., Pilgrim, D., & Nezlin, N. P. (2002). Sea-surface signatures of the island mass effect phenomena around Madeira Island, Northeast Atlantic. Remote Sensing of Environment, 80, 336–360. 10.1016/s0034-4257(01)00316-9

Cañadas, A., & Hammond, P. S. (2008). Abundance and habitat preferences of the short-beaked common dolphin *Delphinus delphis* in the southwestern Mediterranean: Implications for conservation. Endangered Species Research, 4(3), 309–331. 10.3354/esr00073

Cañadas, A., Sagarminaga, R., & García-Tiscar, S. (2002). Cetacean distribution related with depth and slope in the Mediterranean waters off southern Spain. Deep-Sea Research I, 49, 2053–2073. 10.1016/s0967-0637(02)00123-1

Chícaro, M. A., Esteves, E., Santos, A. M. P., dos Santos, A., Peliz, Á., & Ré, P. (2003). Are sardine larvae caught off northern Portugal in winter starving? An approach examining nutritional conditions. Marine Ecology Progress Series, 257, 303–309. 10.3354/meps257303

Correia, A. M., Gil, Á., Valente, R. F., Rosso, M., Sousa-Pinto, I., & Pierce, G. J. (2020). Distribution of cetacean species at a large scale - Connecting continents with the Macaronesian archipelagos in the eastern North Atlantic. Diversity and Distributions, 26(10), 1234–1247. 10.1111/ddi.13127

Correia AM, Gil Á, Valente R, Rosso M, Pierce GJ, Sousa-Pinto I (2019b). Distribution and habitat modelling of common dolphins (*Delphinus delphis*) in the eastern North Atlantic. Journal of the Marine Biological Association of the United Kingdom, 99, 1443–1457. 10.1017/S0025315419000249

Correia, A. M., Sousa-Guedes, D., Gil, Á., Valente, R., Rosso, M., Sousa-Pinto, I., Sillero, N., & Pierce, G. J. (2021). Predicting Cetacean Distributions in the Eastern North Atlantic to Support Marine Management. Frontiers in Marine Science, 8. 10.3389/fmars.2021.643569

Correia, A. M., Tepsich, P., Rosso, M., Caldeira, R., & Sousa-Pinto, I. (2015). Cetacean occurrence and spatial distribution: Habitat modelling for offshore waters in the Portuguese EEZ (NE Atlantic). Journal of Marine Systems, 143, 73–85. 10.1016/j.jmarsys.2014.10.016

Cruz, M. J., Machete, M., Menezes, G., Rogan, E., & Silva, M. A. (2018). Estimating common dolphin bycatch in the pole-and-line tuna fishery in the Azores. PeerJ, 6(2). 10.7717/peerj.4285

Cruz, M. J., Menezes, G., Machete, M., & Silva, M. A. (2016). Predicting interactions between common dolphins and the pole-and-line tuna fishery in the Azores. PLoS ONE, 11(11). 10.1371/journal.pone.0164107

Dawson, S. M., Northridge, S., Waples, D., & Read, A. J. (2013). To ping or not to ping: The use of active acoustic devices in mitigating interactions between small cetaceans and gillnet fisheries. In Endangered Species Research (Vol. 19, Number 3, pp. 201–221). Inter-Research. 10.3354/esr00464

Dias, I. C., Marçalo, A., Feijó, D., Domingos, I., & Silva, A. A. (2022). Interactions between the common dolphin, Delphinus delphis, and the Portuguese purse seine fishery over a period of 15 years (2003–2018). Aquatic Conservation: Marine and Freshwater Ecosystems, 32(8), 1351–1364. 10.1002/aqc.3828

Evans, P. G. H., & Hammond, P. S. (2004). Monitoring cetaceans in European waters. Mammal Rev, 34(1), 131–156. 10.1046/j.0305-1838.2003.00027.x

Fader, J. E., Elliott, B. W., & Read, A. J. (2021). The Challenges of Managing Depredation and Bycatch of Toothed Whales in Pelagic Longline Fisheries: Two U.S. Case Studies. Frontiers in Marine Science, 8. 10.3389/fmars.2021.618031

Feijó, D., Marçalo, A., Bento, T., Barra, J., Marujo, D., Correia, M., & Silva, A. (2018). Trends in the activity pattern, fishing yields, catch and landing composition between 2009 and 2013 from onboard observations in the Portuguese purse seine fleet. Regional Studies in Marine Science, 23, 97–106. 10.1016/j.rsma.2017.12.007

Feijó, D. O. (2013). *Caracterização da pesca do Cerco na Costa Portuguesa*.

Fernández, R., MacLeod, C. D., Pierce, G. J., Covelo, P., López, A., Torres-Palenzuela, J., Valavanis, V., & Santos, M. B. (2013). Inter-specific and seasonal comparison of the niches occupied by small cetaceans off north-west Iberia. Continental Shelf Research, 64, 88–98. 10.1016/j.csr.2013.05.008

Ferreira, M., Eira, C., López, A., & Sequeira, M. (2023). *Delphinus delphis golfinho-comum*.

Garagouni, M., Avgerinou, G., Mouchlianitis, F. A., Minos, G., & Ganias, K. (2022). Questionnaire and experimental surveys show that dolphins cause substantial losses to a gillnet fishery in the eastern Mediterranean Sea. ICES Journal of Marine Science, 79(9), 2552–2561. 10.1093/icesjms/fsac196

Gil, Á. (2017). Cetáceos na Zona Económica Exclusiva Continental Portuguesa: distribuição espaço-temporal e registo de novas ocorrências. Master’s thesis. Universidade do Porto.

Gilles, A., Authier, M., Ramirez-Martinez, N. C., Araújo, H., Blanchard, A., Carlström, J., Eira, C., Dorémus, G., Fernández-Maldonado, C., Geelhoed, S., Kyhn, L., Laran, S., Nachtsheim, D., Panigada, S., Pigeault, R., Sequeira, M., Sveegaard, S., Taylor, N. L., Owen, K., … Hammond, P. S. (2023). Estimates of cetacean abundance in European Atlantic waters in summer 2022 from the SCANS-IV aerial and shipboard surveys. https://www.tiho-hannover.de/itaw/scans-iv-survey

Goetz, S., Read, F. L., Ferreira, M., Portela, J. M., Santos, M. B., Vingada, J., Siebert, U., Marçalo, A., Santos, J., Araújo, H., Monteiro, S., Caldas, M., Riera, M., & Pierce, G. J. (2014). Cetacean occurrence, habitat preferences and potential for cetacean-fishery interactions in Iberian Atlantic waters: Results from cooperative research involving local stakeholders. Aquatic Conservation: Marine and Freshwater Ecosystems, 25(1), 138–154. 10.1002/aqc.2481

Halicka, Z. (2016). Temporal distribution of the short-beaked common dolphin (Delphinus delphis) in the south of Madeira Island (Portugal) and relationship with oceanographic variables. Master’s thesis. Universidade do Algarve.

Hastie, T., & Tibshirani, R. (1986). Generalized Additive Models. Statistical Science, 1(3), 297–318. 10.1214/ss/1177013604

Hintzen, N. T., Brigden, K., Kaastra, H. J., Mackinson, S., Pastoors, M. A., & van de Pol, L. (2025). Bias in Global Fishing Watch AIS data analyses results in overestimate of Northeast Atlantic pelagic fishing impact. ICES Journal of Marine Science, 82(3). 10.1093/icesjms/fsaf033

Kiszka, J. J., Woodstock, M. S., & Heithaus, M. R. (2022). Functional Roles and Ecological Importance of Small Cetaceans in Aquatic Ecosystems. Frontiers in Marine Science, 9. 10.3389/fmars.2022.803173

Lauriano, G., Caramanna, L., Scarnó, M., & Andaloro, F. (2009). An overview of dolphin depredation in Italian artisanal fisheries. Journal of the Marine Biological Association of the United Kingdom, 89(5), 921–929. 10.1017/S0025315409000393

Marçalo, A., Carvalho, F., Frade, M., Bentes, L., Monteiro, P., Pontes, J., Alexandre, S., Oliveira, F., Kingston, A., Erzini, K., & Gonçalves, J. M. S. (2025). Reducing Cetacean Interactions With Bottom Set-Nets and Purse Seining Using Acoustic Deterrent Devices in Southern Iberia. Aquatic Conservation: Marine and Freshwater Ecosystems, 35(2). 10.1002/aqc.70061

Marçalo, A., Katara, I., Feijo, D., Araujo, H., Oliveira, I., Santos, J., Ferreira, M., Monteiro, S., Pierce, G. J., Silva, A., & Vingada, J. (2015). Quantification of interactions between the Portuguese sardine purse-seine fishery and cetaceans. ICES Journal of Marine Science, 72(8), 2438–2449. 10.1093/icesjms/fsv076

Marçalo, A., Nicolau, L., Giménez, J., Ferreira, M., Santos, J., Araújo, H., Silva, A., Vingada, J., & Pierce, G. J. (2018). Feeding ecology of the common dolphin (*Delphinus delphis*) in Western Iberian waters: has the decline in sardine (*Sardina pilchardus*) affected dolphin diet? Marine Biology, 165(3). 10.1007/s00227-018-3285-3

Marçalo, A., Samel, V., Carvalho, F., Frade, M., Erzini, K., & Gonçalves, J. M. (2024). Evaluating dolphin interactions with bottom-set net fisheries off Southern Iberian Atlantic waters. Fisheries Research, 278. 10.1016/j.fishres.2024.107100

Marubini, F., Gimona, A., Evans, P. G. H., Wright, P. J., & Pierce, G. J. (2009). Habitat Preferences and Interannual Variability in Occurrence of the Harbour Porpoise *Phocoena phocoena* off Northwest Scotland. Marine Ecology Progress Series, 381, 297–310. 10.3354/meps07893

Mason, E. (2009). *High-resolution modelling of the Canary Basin oceanic circulation*.

Mendes C, Costa-Dias SC, Teixeira C, Afonso A and Bordalo A (2019). An overview of small scale fisheries in the Northern Portuguese coast. Front. Mar. Sci. Conference Abstract: XX Iberian Symposium on 10.3389/conf.fmars.2019.08.00109

Monteiro, S. S., Torres-Pereira, A., Ferreira, M., Vingada, J. V., Nicolau, L., Sequeira, M., López, A., Covelo, P., Azevedo, M. I., Hernandez-Milian, G., Pierce, G. J., & Eira, C. (2025). What’s on the menu? Diet of common minke whales (*Balaenoptera acutorostrata*) stranded on the Atlantic Iberian coast. Marine Environmental Research, 205. 10.1016/j.marenvres.2025.107024

Moore, J. E., Heinemann, D., Francis, T. B., Hammond, P. S., Long, K. J., Punt, A. E., Reeves, R. R., Sepúlveda, M., Sigurðsson, G. M., Siple, M. C., Víkingsson, G. A., Wade, P. R., Williams, R., & Zerbini, A. N. (2021). Estimating Bycatch Mortality for Marine Mammals: Concepts and Best Practices. In Frontiers in Marine Science (Vol. 8). Frontiers Media S.A. 10.3389/fmars.2021.752356

Moura, A. E., Sillero, N., & Rodrigues, A. (2012). Common dolphin (*Delphinus delphis*) habitat preferences using data from two platforms of opportunity. Acta Oecologica, 38, 24–32. 10.1016/j.actao.2011.08.006

Paradell, O. G., López, B. D., & Methion, S. (2019). Modelling common dolphin (*Delphinus delphis*)coastal distribution and habitat use: Insights for conservation. Ocean and Coastal Management, 179. 10.1016/j.ocecoaman.2019.104836

Pearce, J. L., & Boyce, M. S. (2006). Modelling distribution and abundance with presence-only data. In Journal of Applied Ecology (Vol. 43, Number 3, pp. 405–412). 10.1111/j.1365-2664.2005.01112.x

Powell, J. R., & Wells, R. S. (2011). Recreational fishing depredation and associated behaviors involving common bottlenose dolphins (*Tursiops truncatus*) in Sarasota Bay, Florida. Marine Mammal Science, 27(1), 111–129. 10.1111/j.1748-7692.2010.00401.x

Ré, P., Silva, R. C. e, Cunha, E., Farinha, A., Meneses, I., & Moita, T. (1990). Sardine spawning off Portugal. https://www.researchgate.net/publication/285107053

Read, A. J. (2008). The looming crisis: interactions between marine mammals and fisheries. Journal of Mammalogy, 89(3), 541–548. 10.1644/07-mamm-s-315r1.1

Roberts, J. J., Best, B. D., Dunn, D. C., Treml, E. A., & Halpin, P. N. (2010). Marine Geospatial Ecology Tools: An integrated framework for ecological geoprocessing with ArcGIS, Python, R, MATLAB, and C++. Environmental Modelling and Software, 25(10), 1197–1207. 10.1016/j.envsoft.2010.03.029

Rufino, M. M., Mendo, T., Samarão, J., & Gaspar, M. B. (2023). Estimating fishing effort in small-scale fisheries using high-resolution spatio-temporal tracking data (an implementation framework illustrated with case studies from Portugal). Ecological Indicators, 154. 10.1016/j.ecolind.2023.110628

Santos, M. B., González-Quirós, R., Riveiro, I., Cabanas, J. M., Porteiro, C., & Pierce, G. J. (2012). Cycles, trends, and residual variation in the Iberian sardine (*Sardina pilchardus*) recruitment series and their relationship with the environment. ICES Journal of Marine Science, 69(5), 739–750. 10.1093/icesjms/fsr186

Santos, M. B., Saavedra, C., & Pierce, G. J. (2014). Quantifying the predation on sardine and hake by cetaceans in the Atlantic waters of the Iberian Peninsula. Deep-Sea Research Part II: Topical Studies in Oceanography, 106, 232–244. 10.1016/j.dsr2.2013.09.040

Silva, F. L. da, Grassi Sella, M. L., Francoy, T. M., & Costa, A. H. R. (2015). Evaluating classification and feature selection techniques for honeybee subspecies identification using wing images. Computers and Electronics in Agriculture, 114, 68–77. 10.1016/j.compag.2015.03.012

Silva, M. A. (1999). Diet of common dolphins, *Delphinus delphis*, off the Portuguese continental coast. Journal of the Marine Biological Association of the United Kingdom, 79(3), 531–540. 10.1017/S0025315498000654

Silva, M. A., & Sequeira, M. (2003). Patterns in the mortality of common dolphins (Delphinus delphis) on the Portuguese coast, using stranding data records, 1975-1998. Aquatic Mammals, 29, 88–98. 10.1578/016754203101023924

Sousa, P., Azevedo, M., & Gomes, M. C. (2005). Demersal assemblages off Portugal: Mapping, seasonal, and temporal patterns. Fisheries Research, 75(1–3), 120–137. 10.1016/j.fishres.2005.03.012

Spadoni, G., Duarte, R., Soares, C., Fernandez, M., & Jesus, S. M. (2025). Common dolphin’s shipping noise risk assessment on the Portuguese coast. Marine Pollution Bulletin, 211. 10.1016/j.marpolbul.2024.117415

Tack, K. (2025). Toward a Global Estimate of Cetacean Bycatch: Data Gaps and Future Needs.

Temple, A. J., Langner, U., & Berumen, M. L. (2024). Management and research efforts are failing dolphins, porpoises, and other toothed whales. Scientific Reports, 14(1). 10.1038/s41598-024-57811-7

Veiga-Malta, T., Szalaj, D., Angélico, M. M., Azevedo, M., Farias, I., Garrido, S., Lourenço, S., Marçalo, A., Marques, V., Moreno, A., Oliveira, P. B., Paiva, V. H., Prista, N., Silva, C., Sobrinho-Gonçalves, L., Vingada, J., & Silva, A. (2019). First representation of the trophic structure and functioning of the Portuguese continental shelf ecosystem: Insights into the role of sardine. Marine Ecology Progress Series, 617–618, 323–340. 10.3354/meps12724

Vingada, J.V., Eira, C. (2018). Conservação de Cetáceos e Aves Marinhas em Portugal Continental. O projeto LIFE+ MarPro. LIFE. Rainho & Neves, Lda., Santa Maria da Feira.

Vitorino, J., Oliveira, A., Jouanneau, J. M., & Drago, T. (2002). Winter dynamics on the northern Portuguese shelf. Part 1: physical processes. Progress in Oceanography, 52, 129–153. 10.1016/s0079-6611(02)00003-4

Wardhani, Lita T A L, Herawati, R., Pinilih, S. A. G., Ristyawati, A., & Wardhani, L T A L. (2022). Global Fishing Watch System as a solution in the control of the fishing industry in Indonesia (Vol. 15). http://www.bioflux.com.ro/aacl

Wise, L., Silva, A. A., Ferreira, M., & Silva, M. A. (2007). Interactions between small cetaceans and the purse-seine fishery in western Portuguese waters. Scientia Marina.

Wood, S. N. (2003). Thin plate regression splines. J. R. Statist. Soc. B, 65(1), 95–114. 10.1111/1467-9868.00374

Wood, S. N. (2004). Stable and efficient multiple smoothing parameter estimation for generalized additive models. Journal of the American Statistical Association, 99(467), 673–686. 10.1198/016214504000000980

Wood, S. N. (2011). Fast stable restricted maximum likelihood and marginal likelihood estimation of semiparametric generalized linear models. J. R. Statist. Soc. B, 73(1), 3–36. 10.1111/j.1467-9868.2010.00749.x

Zuur, A. F., Ieno, E. N., & Elphick, C. S. (2010). A protocol for data exploration to avoid common statistical problems. Methods in Ecology and Evolution, 1(1), 3–14. 10.1111/j.2041-210x.2009.00001.x

Zuur, A. F., Ieno, E. N., & Smith, G. M. (2007). Analysing Ecological Data. In Series Editors (Ed.), Statistics for Biology and Health. Springer. 10.1007/978-0-387-45972-1

Zwolinski, J. P., Oliveira, P. B., Quintino, V., & Stratoudakis, Y. (2010). Sardine potential habitat and environmental forcing off western Portugal. ICES Journal of Marine Science, 67, 1553–1564. 10.1093/icesjms/fsq068

