## Supplementary material for "Exploring the relationship between dolphins and fisheries: uncovering the spatial and temporal patterns that influence potential conflicts along Portugal’s north coast"

Table 1. Environmental data with the corresponding spatial and temporal resolutions and datasets.

| Variable | Spatial Resolution | Temporal Resolution | Dataset | Source |
|---|---|---|---|---|
| Bathymetry | 15 arc-second | ------------ | GEBCO_2021<br>GEBCO_2022<br>GEBCO_2023<br>GEBCO_2024 | GEBCO (2021, 2022, 2023, 2024) |
| Sea Surface Temperature (SST) | 0.0083° | Monthly | cmems_mod_glo_phy_my_0.083deg_P1M-m (01/01/1993–01/06/2021)<br>cmems_mod_glo_phy_myint_0.083deg_P1M-m (01/07/2021–01/04/2025) | CMEMS (1993-2025) |
| | | Daily | cmems_mod_glo_phy_my_0.083deg_P1D-m (01/01/1993–30/06/2021)<br>cmems_mod_glo_phy_myint_0.083deg_P1D-m (01/07/2021–22/04/2025) | CMEMS (1993-2025) |
| Chlorophyll-a concentration (CHL) | 4 km | Monthly | cmems_obs-oc_glo_bgc-plankton_my_l4-multi-4km_P1M (01/09/1997–01/06/2025) | CMEMS (1997-2025) |
| | 300 m | | cmems_obs-oc_glo_bgc-plankton_my_l4-olci-300m_P1M (01/04/2016–01/06/2025) | CMEMS (2016-2025) |
| | 4 km | Daily | cmems_obs-oc_glo_bgc-plankton_my_l4-gapfree-multi-4km_P1D (04/09/1997–15/07/2025) | CMEMS (1997-2025) |
| Sea Surface Salinity (SSS) | 0.0083° | Monthly | cmems_mod_glo_phy_my_0.083deg_P1M-m (01/01/1993–01/06/2021)<br>cmems_mod_glo_phy_myint_0.083deg_P1M-m (01/07/2021–01/04/2025) | CMEMS (1993-2025) |
| | | Daily | cmems_mod_glo_phy_my_0.083deg_P1D-m (01/01/1993–30/06/2021)<br>cmems_mod_glo_phy_myint_0.083deg_P1D-m (01/07/2021–22/04/2025) | CMEMS (1993-2025) |

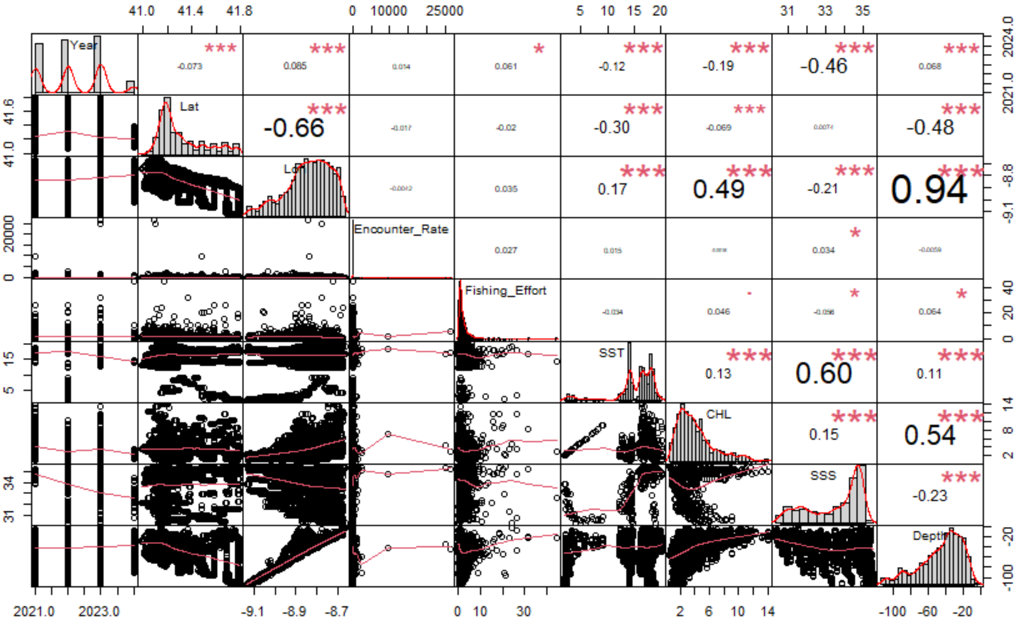


Figure 1. Pearson correlation matrix between all pairs of explanatory variables.

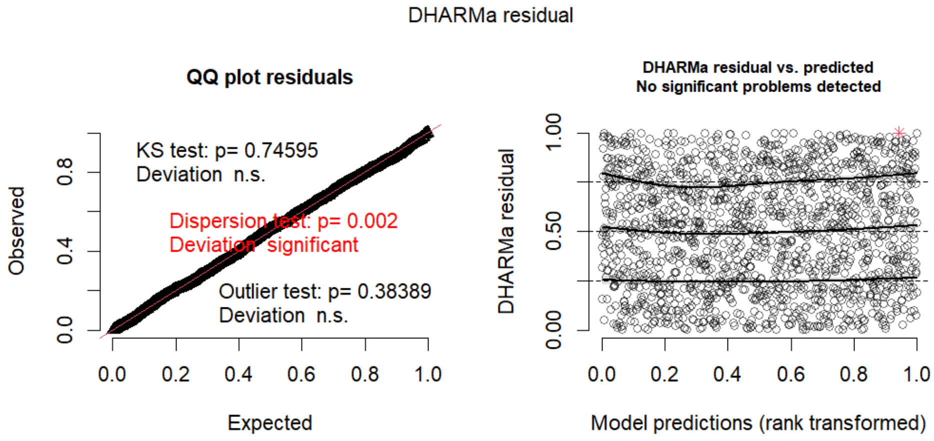


Figure 2. DHARMa residual diagnostics for the fitted GAM.

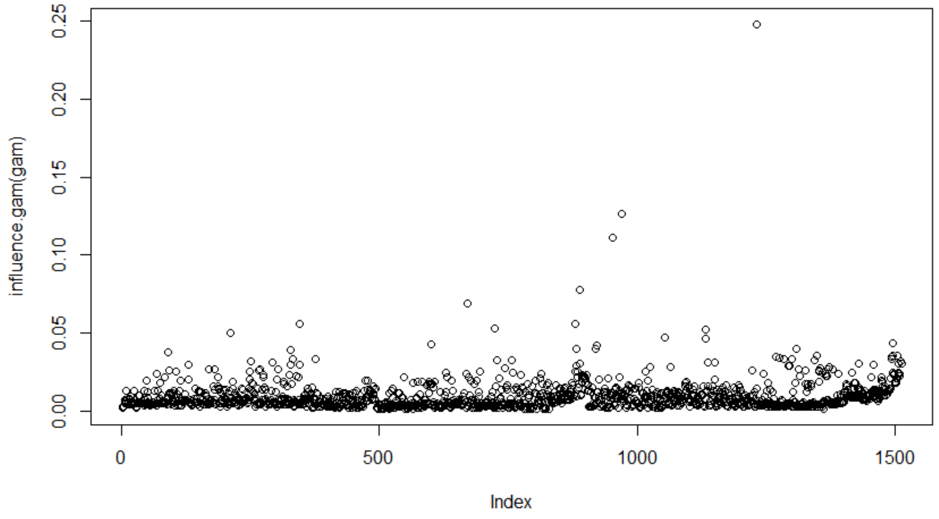


Figure 3. Influence data points of the best GAM.

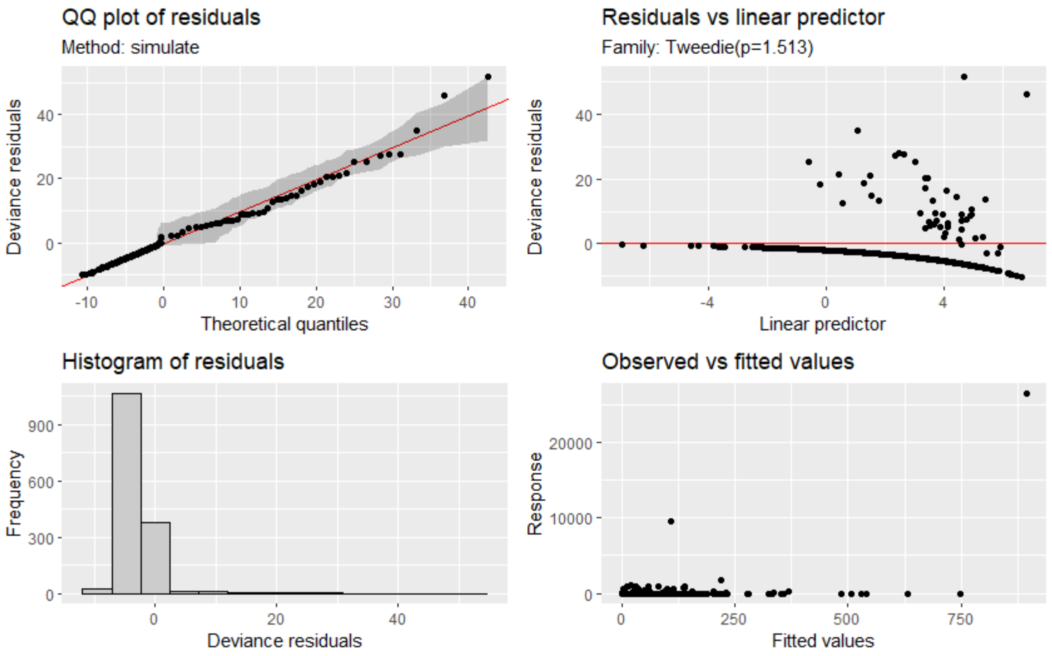


Figure 4. Diagnostic plots produced by "appraise ()" function (gratia package) for the fitted GAM, including QQ-plot of residuals, residuals versus linear predictor, histogram of residuals, and observed versus fitted values.
